# Speech-related auditory salience detection in the posterior superior temporal region

**DOI:** 10.1101/2021.04.14.439910

**Authors:** Erik C. Brown, Brittany Stedelin, Ahmed M. Raslan, Nathan R. Selden

## Abstract

Processing auditory human speech requires both detection (early and transient) and analysis (sustained). We analyzed high gamma (70-110Hz) activity of intracranial electroencephalography waveforms acquired during an auditory task that paired forward speech, reverse speech, and signal correlated noise. We identified widespread superior temporal sites with sustained activity responding only to forward and reverse speech regardless of paired order. More localized superior temporal auditory onset sites responded to all stimulus types when presented first in a pair and responded in recurrent fashion to the second paired stimulus in select conditions even in the absence of interstimulus silence; a novel finding. Auditory onset activity to a second paired sound recurred according to relative salience, with evidence of partial suppression during linguistic processing. We propose that temporal lobe auditory onset sites facilitate a salience detector function with hysteresis of 200ms and are influenced by cortico-cortical feedback loops involving linguistic processing and articulation.

## Introduction

Since Wernicke and Lichtheim first conceived their model of language structure and function,^1^ a localizationist view of brain function has persisted^2^. Historically, Wernicke’s region has been functionally defined as a language comprehension center, lacking clear anatomical boundaries^3^. Although the posterior portion of the superior temporal gyrus (pSTG) is the anatomical site most commonly assigned to Wernicke’s region^4,5^, some evidence suggests that the pSTG does not have a direct role in language processing^6,7^ or does not participate in language comprehension at all^4^.

Intracranial electroencephalography (iEEG) has yielded unique insights into many aspects of brain function, including language. Brown et al. described two separable functional auditory language sites; ‘Early Auditory’ and ‘Full Auditory’^6,7^. In contrast to Full Auditory sites, Early Auditory sites exhibited only brief, early high gamma range augmentation following auditory stimulus onset and responded to non-speech noise in the same way as they responded to speech. Hamilton et al. described the same phenomenon, using the terms ‘Sustained’ and ‘Onset’, describing Onset sites to be non-discriminatory and consistent in both time and space^8^. Forseth et al. also detailed this phenomenon, relating ‘Transient’ sites with stimulation induced speech production deficit^9^. These previous findings correspond in localizing ‘Onset/Transient/Early Auditory’ sites to more posterior superior temporal locations.

To further elucidate the underlying functional relevance of pSTG sites, we aimed to test the hypothesis of Brown et al. (2014) that Early Auditory sites subserve human voice detection. The terminology ‘Onset’ and ‘Sustained’, as described by Hamilton et al. as well as the larger neuroscientific literature, will be employed throughout^10,11^.

To effectively facilitate language function, a human voice detector might function similarly to a network switchboard deploying a routing protocol. Complex routing protocols enable the function of dense networks in computer science^12^. In neuroscience, higher order relays may mediate corticocortical communication via efference copies^13^ that may be collated with reafferent information for cross-modality cortical control networks^14^. Such a system should be sensitive to afferent signal state changes. Our primary analysis aims to determine if Onset sites are sensitive to sudden state changes in an auditory stimulus. To achieve this we designed a language task to present paired 1s duration stimuli of forward speech, reverse speech, and signal correlated noise (SCN), with no pause between pairs. Assuming that Onset sites are part of an auditory input monitoring network, we hypothesized that high gamma activity (70-110Hz) related to auditory stimulus trial onset would recur when that stimulus suddenly changes state mid-trial.

## Results

### Participants

Seventeen patients (8male, 9 female; aged between 11-57 years; **Table 1**) met the study inclusion criteria. A total of 2,028 electrodes (1,712 depth, 316 subdural) were implanted.

**Table 1.** Patient Data

| Patient | Sex | Age at surgery (years) | Dominant hand | Age at epilepsy onset (years) | Language Lateralization |  | Neuropsychiatric Testing |  |  |  |  | Response Time (s) |  |  | Seizure semiology | Electrodes | Temporal Sites (n) |  |
| --- | --- | --- | --- | --- | --- | --- | --- | --- | --- | --- | --- | --- | --- | --- | --- | --- | --- | --- |
|  |  |  |  |  | fMRI |  | WADA | WTAR (SS) | COWAT (SS) | PSI (SS) | VCI (SS) | mean | median | SD |  | (side, n, location) | Sustained | Onset |
|  |  |  |  |  | Expressive | Receptive |  |  |  |  |  |  |  |  |  |  |  |  |
| 1 | F | 13 | Rt | 7 | Lt | Undefined | N/A | N/A | N/A | 86 | 89 | 3.11 | 3.01 | 0.56 | Focal | Rt 88 subdural | 3 | 1 |
| 2 | M | 11 | Rt | 2 | N/A | N/A | N/A | N/A | N/A | 83 | 103 | 2.85 | 2.92 | 0.22 | Focal | Rt 144 depth | 0 | 0 |
| 3 | M | 39 | Rt | 20 | Lt | Lt | N/A | 88 | N/A | N/A | N/A | 2.97 | 3.04 | 0.39 | Focal | Lt 98 depth, 32 subdural | 2 | 0 |
| 4 | M | 33 | Lt | 23 | Lt>Rt | Lt>Rt | N/A | N/A | N/A | 100 | 125 | 2.92 | 3.08 | 0.30 | Focal | Lt 88 depth, 28 subdural; Rt 10 depth | 4 Lt | 0 |
| 5 | F | 16 | Rt | 2 | Rt>Lt | Rt | N/A | N/A | N/A | 92 | 80 | 4.11 | 3.62 | 1.88 | Focal | Rt 80 subdural, 22 depth | 6 | 5 |
| 6 | F | 12 | Rt | 1 | Lt | Lt | N/A | N/A | N/A | 105 | 86 | 11.14 | 5.68 | 9.84 | Focal | Lt 64 depth; Rt 20 depth | 7 Lt, 1 Rt | 0 |
| 7 | F | 57 | Rt | 50 | Lt | Lt | N/A | 96 | 95 | N/A | N/A | 4.32 | 3.92 | 1.24 | 2° General | Lt 55 depth; Rt 70 depth | 4 Rt | 1 Lt |
| 8 | M | 39 | Rt | 36 | Lt | Lt | N/A | 86 | N/A | N/A | 80 | 3.32 | 3.04 | 0.73 | 2° General | Lt 118 depth; Rt 8 depth | 8 Lt | 2 Lt |
| 9 | F | 31 | Rt | 24 | Lt | Lt | N/A | 119 | N/A | N/A | 106 | 3.19 | 3.23 | 0.28 | Focal | Lt 106 depth; Rt 20 depth | 8 Lt, 2 Rt | 1 Lt |
| 10 | M | 30 | Lt | 17 | Lt | Undefined | Lt | 108 | N/A | N/A | N/A | 3.61 | 3.26 | 0.71 | Focal | Lt 82 depth; Rt 46 depth | 11 Lt, 3 Rt | 5 Lt, 3 Rt |
| 11 | F | 33 | Rt | 22 | Lt | Lt | Lt | N/A | N/A | 94 | 91 | 3.51 | 3.54 | 0.52 | Focal | Lt 96 depth; Rt 32 depth | 5 Lt | 0 |
| 12 | M | 25 | Rt | 21 | Lt | Lt | Lt | N/A | 94 | N/A | N/A | 3.42 | 3.34 | 0.51 | 2° General | Lt 48 depth; Rt 80 depth | 2 Lt, 6 Rt | 0 |
| 13 | F | 38 | Rt | 12 | Lt | Lt | N/A | 112 | 74 | N/A | N/A | 2.74 | 2.64 | 0.35 | Focal | Lt 79 depth; Rt 48 depth | 1 Lt, 2 Rt | 0 |
| 14 | M | 11 | Rt | 11 | Lt | Lt | N/A | N/A | N/A | N/A | 103 | 2.99 | 2.76 | 0.54 | 2° General | Lt 88 subdural | 2 | 1 |
| 15 | F | 20 | Rt | 16 | Lt | Lt | Lt | N/A | N/A | 127 | 107 | 2.71 | 2.68 | 0.34 | 2° General | Lt 98 depth; Rt 30 depth | 3 Lt | 3 Lt |
| 16 | M | 37 | Rt | 35 | Lt | Lt | Lt | N/A | N/A | 97 | 105 | 3.46 | 3.43 | 0.60 | Focal | Lt 104 depth; Rt 24 depth | 14 Lt, 3 Rt | 2 Lt |
| 17 | F | 31 | Rt | 12 | Lt & Rt | Undefined | N/A | N/A | N/A | N/A | N/A | 3.88 | 3.76 | 1.19 | Focal | Lt 22 depth; Rt 100 depth | 3 Rt | 0 |
| F: Female. M: Male. Lt: Left. Rt: Right. fMRI; functional magnetic resonance imaging. WTAR: Wechsler Test of Adult Reading. COWAT: Controlled Oral Word Association Test. PSI: Processing Speed Index. VCI: Verbal Comprehension Index. N/A: Not Applicable. SS: Standard Score. SD: Standard Deviation |  |  |  |  |  |  |  |  |  |  |  |  |  |  | Total (n, side) | 1206 Lt, 822 Rt | 67 Lt, 33 Rt | 14 Lt, 10 Rt |

### Hemispheric Dominance and Behavioral Data

All 17 patients completed the language task, remaining awake and attentive throughout. Hemispheric language dominance was evaluated in 16 of the 17 patients with pre-operative clinical functional magnetic resonance imaging (fMRI). Of these 16 patients, 13 were confirmed to be left-hemisphere dominant for expressive language and 11 for receptive language. In patients 1, 10, and 17, receptive language function could not be delineated in either hemisphere as assessed by fMRI; patient 17 was codominant for expressive language. Patient 4 exhibited evidence for asymmetric codominance of both expressive and receptive language function, favoring the left side. Patient 5 was the only patient with evidence of right hemisphere language dominance as assessed by fMRI. Patients 10-12, 15, and 16 all underwent Wada testing; all with left hemisphere language dominance. Patient clinical characteristics, hemisphere dominance, and response times for correct responses across all English language question trials, measured from stimulus onset to response onset, are summarized in **Table 1**. Across all patients and trials, only one question answered by patient 1, was not answered correctly. Patient 3 heard the first question stimulus but did not immediately answer; after a reminder of the instructions the patient performed well. Patient 6 had greatly increased response time to three of the question stimuli not representative of overall performance, as observed by separation of mean and median. Although all three were answered correctly, these three trials were later excluded from response-onset iEEG analysis. Patient 17 answered every English language question correctly, responding to only one in Spanish. This bilingual patient also received the same questions translated to Spanish as spoken by a non-native speaker. They had difficulty understanding two of the Spanish language versions of the questions, answering the remaining seven with the following parameters for comparison: mean 4.11s, median 4.36s, and standard deviation 1.55s.

### Cortical Response of Sustained Temporal Auditory Sites to Speech Trial Pairs

Based upon cortical responses to randomly presented English language auditory questions, we identified a total of 100 electrode contacts within the temporal lobe classified as Auditory that were subclassified as Sustained. Cohort level findings are detailed in **Table 2** and representative site results from Patient 5 in **Fig. 1**.

**Table 2.**
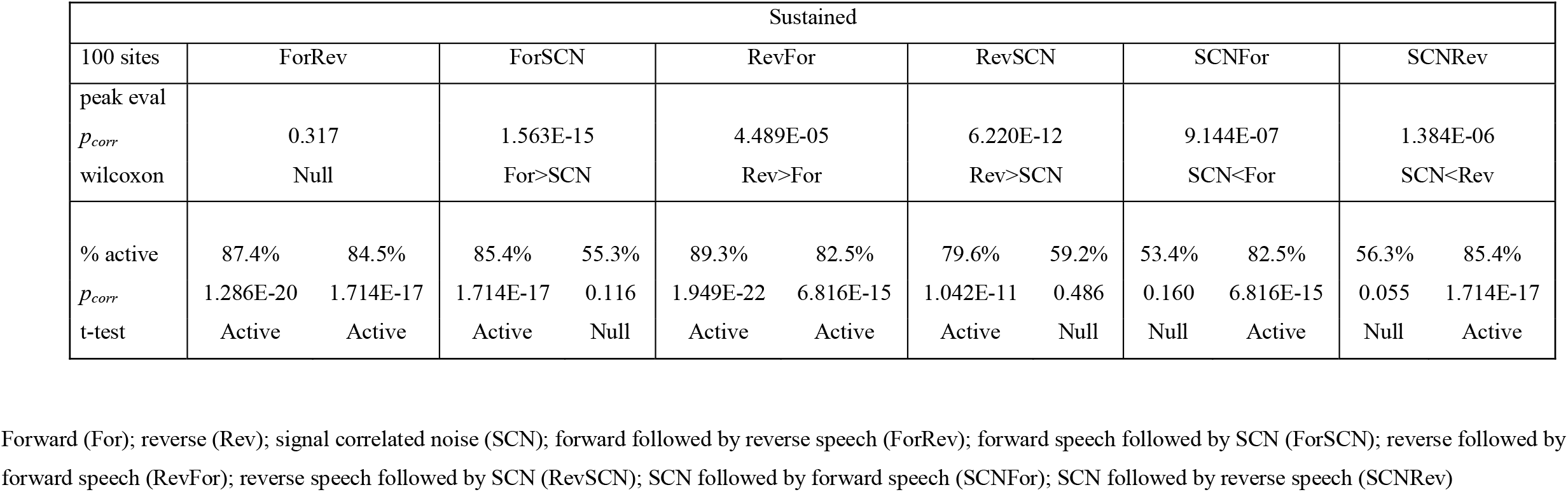
Cohort Statistics – Sustained Sites

**Fig. 1.**
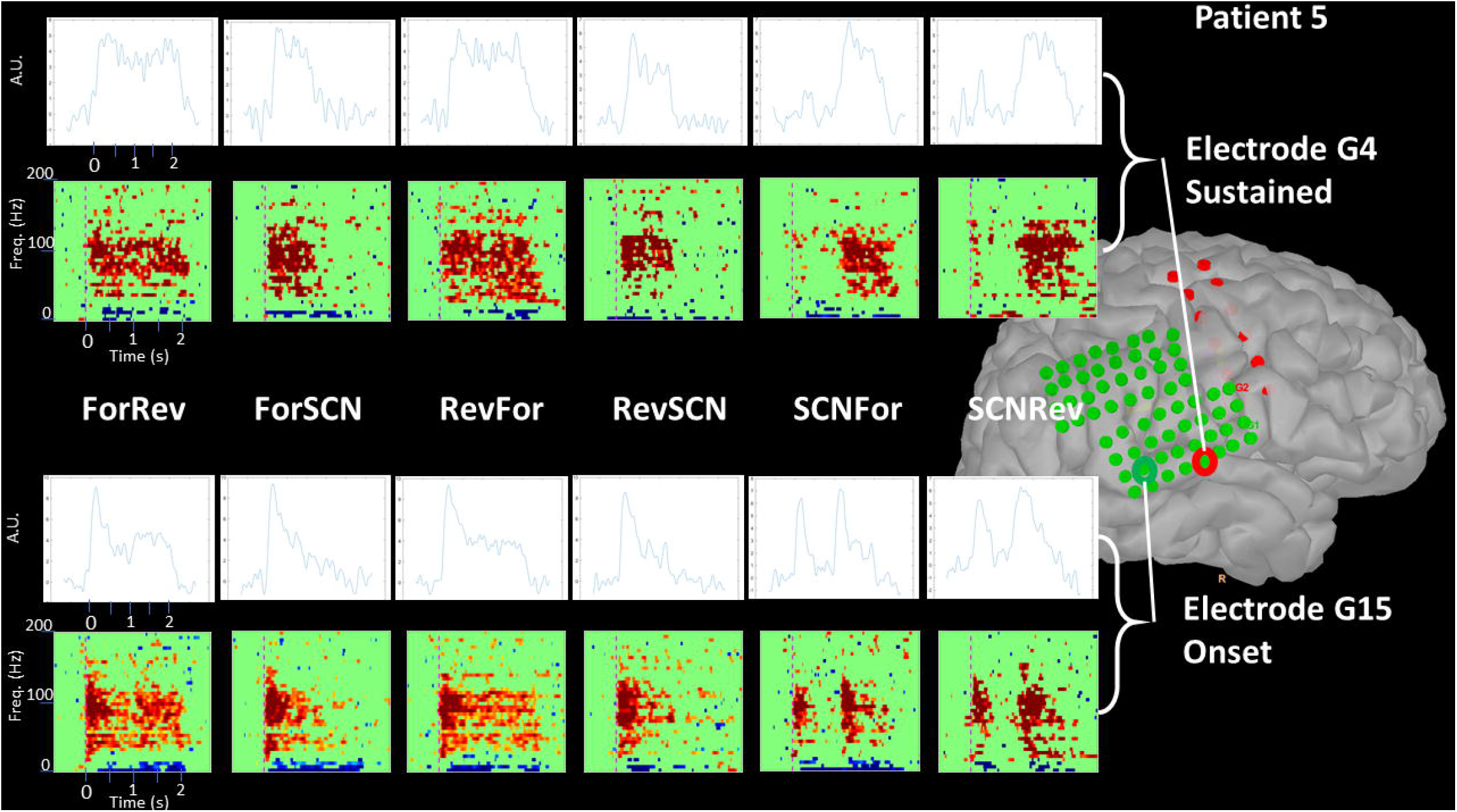
Cortical Response at Select Temporal Onset and Sustained Sites to Paired Stimuli. Displayed response profiles from a representative Onset (bottom) and Sustained (top) sites from patient 5. The upper row of graphs for each site show profiles of average high gamma activity across 70 to 110Hz with y-axis of Arbitrary Units (A.U.) representing values relative to baseline and x-axis representing time in milliseconds (ms). The lower row of graph for each site are statistical frequency (Hz) by time (s) plots with red representing significant augmentation and blue representing significant attenuation relative to baseline. The brain inset to the right depicts the location of the selected electrodes on this patient’s right hemisphere, which was their language dominant hemisphere.

When forward speech was presented first in a pair, significant high gamma augmentation was observed more often than chance in both the forward followed by reverse speech (ForRev) and the forward speech followed by SCN (ForSCN) conditions, 87% (p_corr_ = 1.29×10^−20^) and 85% (p_corr_ = 1.71×10^− 17^), respectively. When Reverse speech was presented first in a pair, significant high gamma augmentation was observed more often than chance in both the reverse followed by forward speech (RevFor) and the reverse speech followed by SCN (RevSCN) conditions, 89% (p_corr_ = 1.95×10^−22^) and 80% (p_corr_ = 1.04×10^− 11^), respectively. When SCN was presented first in a pair, significant high gamma augmentation was not observed more often than chance in either the SCN followed by forward speech (SCNFor) or the SCN followed by reverse speech (SCNRev) conditions, 53% (p = 0.160) and 56% (p = 0.055), respectively.

When forward speech was presented second in a pair, significant high gamma augmentation was observed more often than chance in both the RevFor and the SCNFor conditions, 83% (p_corr_ = 6.82×10^−15^) and 83% (p_corr_ = 6.82×10^−15^), respectively. When Reverse speech was presented second in a pair, significant high gamma augmentation was observed more often than chance in both the ForRev and the SCNRev conditions, 84% (p_corr_ = 1.71×10^−17^) and 85% (p_corr_ = 1.71×10^−17^), respectively. When SCN was presented second in a pair, significant high gamma augmentation was not observed more often than chance in either the ForSCN or the RevSCN conditions, 55% (p = 0.116) and 59% (p_corr_ = 0.486), respectively.

Unique to the ForRev condition, peak high gamma event-related spectral perturbation (ERSP) values did not differ between the first and second sounds of the pair (p = 0.317). Peak ERSP during forward speech was higher than that during SCN in both ForSCN (p_corr_ = 1.56×10^−15^) and SCNFor (p_corr_ = 9.14×10^−7^) conditions. Peak ERSP during reverse speech was higher than that during SCN in both the RevSCN (p_corr_ = 6.22×10^−12^) and SCNRev (p_corr_ = 1.38×10^−6^) conditions. Peak ERSP during reverse speech was higher than that during forward speech in the RevFor condition (p_corr_ = 4.49×10^−5^).

### Cortical Response of Onset Temporal Auditory Sites to Speech Trial Pairs

Based upon cortical responses to randomly presented English language auditory questions, we identified a total of 24 electrode contacts within the temporal lobe classified as Auditory to be subclassified as Onset. Cohort level findings are detailed in **Table 3** and representative site results from Patient 5 in **Fig. 1**.

**Table 3.** Cohort Statistics – Onset Sites

| Onset |  |  |  |  |  |  |  |  |  |  |  |  |
| --- | --- | --- | --- | --- | --- | --- | --- | --- | --- | --- | --- | --- |
| 24 sites | ForRev |  | ForSCN |  | RevFor |  | RevSCN |  | SCNFor |  | SCNRev |  |
| peaks |  |  |  |  |  |  |  |  |  |  |  |  |
| $p_{corr}$ | 1.156 | | 2.280E-03 | | 4.130E-03 | | 6.550E-03 | | 0.162 | | 0.668 | |
| wilcoxon | Null |  | For>SCN |  | Rev>For |  | Rev>SCN |  | Null |  | Null |  |
| % active | 95.8% | 91.7% | 95.8% | 66.7% | 95.8% | 79.2% | 95.8% | 79.2% | 83.3% | 87.5% | 87.5% | 91.7% |
| $p_{corr}$ | 2.212E-09 | 4.188E-06 | 2.212E-09 | 0.052 | 2.212E-09 | 3.960E-02 | 2.212E-09 | 3.960E-02 | 4.926E-03 | 2.852E-04 | 2.852E-04 | 4.188E-06 |
| t-test | Active | Active | Active | Null | Active | Active | Active | Active | Active | Active | Active | Active |

Forward (For); reverse (Rev); signal correlated noise (SCN); forward followed by reverse speech (ForRev); forward speech followed by SCN (ForSCN); reverse followed by forward speech (RevFor); reverse speech followed by SCN (RevSCN); SCN followed by forward speech (SCNFor); SCN followed by reverse speech (SCNRev)

In all conditions, regardless of type of sound, the first sound of the pair was associated with statistically significant high gamma augmentation occurring more often than chance. Rates of high gamma augmentation for the first sound in the pair condition were: ForRev 96% (p_corr_ = 2.21×10^−9^), ForSCN 96% (p_corr_ = 2.21×10^−9^), RevFor 96% (p_corr_ = 2.21×10^−9^), RevSCN 96% (p_corr_-= 2.21×10^−9^), SCNFor 83% (p_corr_ = 4.93×10^−3^), and SCNRev 88% (p_corr_ = 2.85×10^−4^).

When forward speech was presented second in a pair, significant high gamma augmentation was observed more often than chance in both the SCNFor and the RevFor conditions, 88% (p_corr_ = 2.85×10^−4^) and 79% (p_corr_ = 0.040), respectively. When reverse speech was presented second in a pair, significant high gamma augmentation was observed more often than chance in both the ForRev and the SCNRev conditions, 92% (p_corr_ = 4.19×10^−6^) and 92% (p_corr_ = 4.19×10^−6^), respectively. When SCN was presented second in a pair, significant high gamma augmentation was observed more often than chance in the RevSCN condition but not in the ForSCN condition, 79% (p_corr_ = 0.040) and 67% (p_corr_ = 0.052), respectively.

In the ForRev, SCNFor, and SCNRev conditions, peak ERSP values did not differ between the first and second sounds of the pair (p_corr_ = 1.16, p = 0.162, and p = 0.668, respectively). Peak ERSP during reverse speech was higher than that during forward in the RevFor (p_corr_ = 4.13×10^−3^) condition. Peak ERSP during SCN was lower than that during the preceding sound in both the ForSCN (p_corr_ = 2.28×10^−3^) and RevSCN (p_corr_ = 6.55×10^−3^) conditions.

### Comparison of Onset and Sustained Auditory Sites of the Temporal Lobe

Temporal lobe sites identified as Sustained are rather widespread through superior aspects of the middle to posterior temporal lobe, both superficially and deep (**Fig. 2** and supplemental **Video S1**). Contrarily, temporal lobe sites identified as Onset clearly favor more posterior aspects of the superior temporal region including the superior temporal gyrus and deep within the superior temporal sulcus with few exceptions in the dominant hemisphere. The more sparsely sampled non-dominant hemisphere possessed a less well-defined spatial pattern. When comparing the latencies between Sustained and Onset sites across the 9 patients that exhibited sites of both types, we find with statistical significance that the median latency values across Sustained sites is longer than that across Onset sites (107ms vs. 82ms, p = 0.0209).

**Fig. 2.**
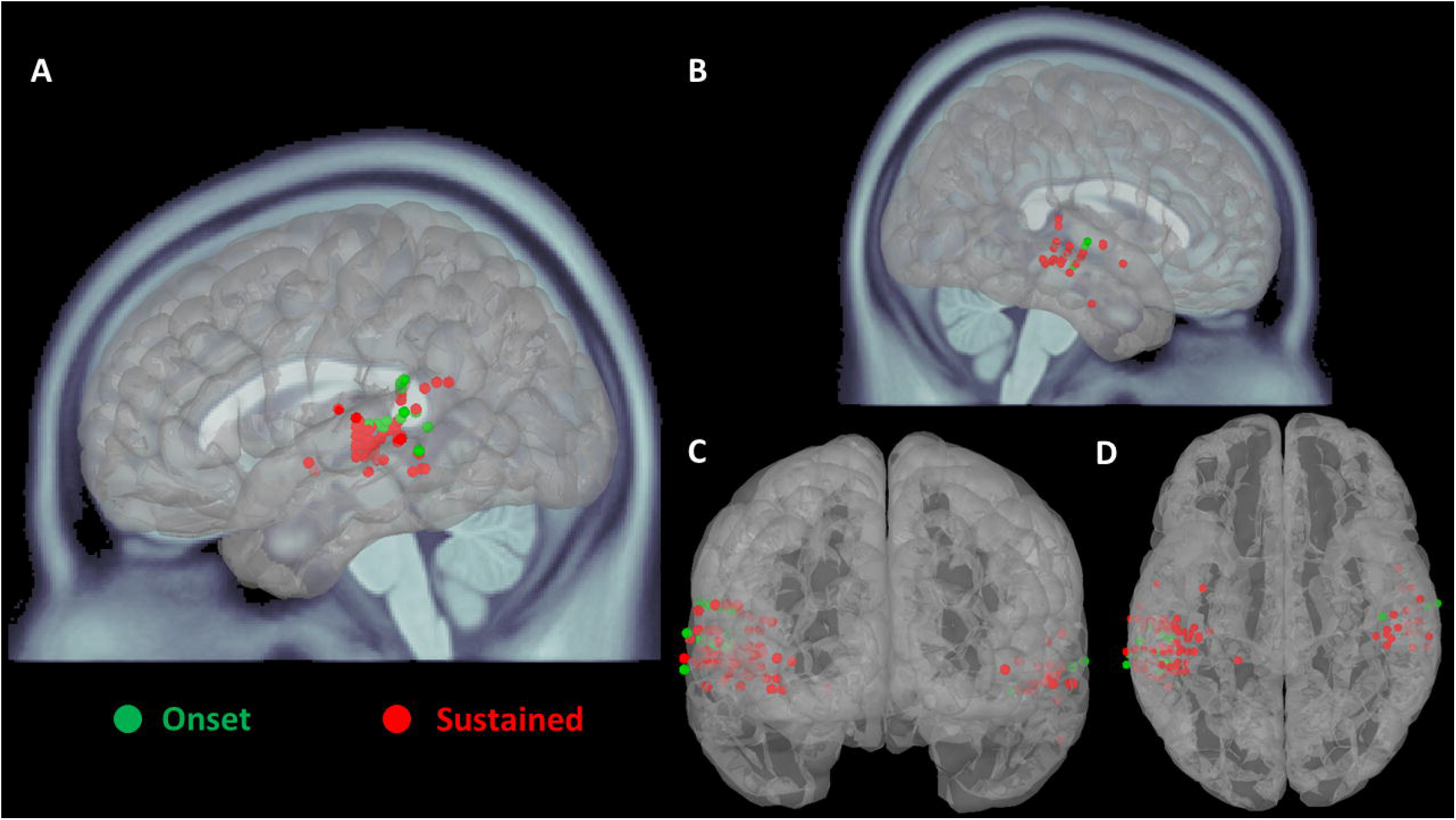
Cohort Level Representation of All Active Temporal Lobe Auditory Sites. Displayed is the cohort level representation of all active temporal lobe auditory sites to visualize localization of Onset and Sustained sites on the ICBM152 atlas brain. (A) and (B) dominant and non-dominant hemispheres, respectively; note the poor sampling of the non-dominant hemisphere. (C) and (D) orient the three-dimension atlas brain in posterior-anterior and superior-inferior perspectives, respectively, in order to appreciate depth of active electrode locations. Green colored electrodes sites represent Onset, and red color represents Sustained.

Sustained sites active during SCN sounds in the SCNFor and SCNRev conditions were 53% and 56% of the time, respectively. Despite lack of evidence that either condition statistically exceeded the 50% chance threshold, this was a surprising finding^7^. Considering previous data^7^ were collected strictly from subdural electrodes, we divided the Sustained sites into ‘Lateral’ and ‘Deep’, with Lateral sites defined as those within 1cm of the lateral pial surface and Deep the remainder, to explore if Lateral Sustained sites behave differently than Deep Sustained sites. With this simple definition, we found 31 of the Sustained sites to be Lateral and 69 to be Deep. For Lateral Sustained sites, we find such activity during SCN when presented first in a stimulus pair to occur at 39% during the SCNFor condition and at 32% during the SCNRev condition. For Deep Sustained sites, this occurred at 62% during the SCNFor condition and at 70% during the SCNRev condition. When combining rates of significant high gamma augmentation during SCN when presented first in a pair, we have insufficient statistical power to show this to occur less often than chance for Lateral Sustained sites, 35% (p_corr_ = 0.798). However, it occurs more often than chance for Deep Sustained sites, 66% (p_corr_ = 4.96×10^−3^). This was detected per two-tailed t-test comparing the rate to 50% chance with Bonferroni correction for 38 statistical comparisons of high gamma-augmentation during paired sound stimuli. Qualitative inspection reveals that many Deep Sustained sites appear to possess elements of both Onset and Lateral Sustained sites (**Fig. 3** from patient 9 who expressed Lateral Sustained, Deep Sustained, and Onset sites).

**Fig. 3.**
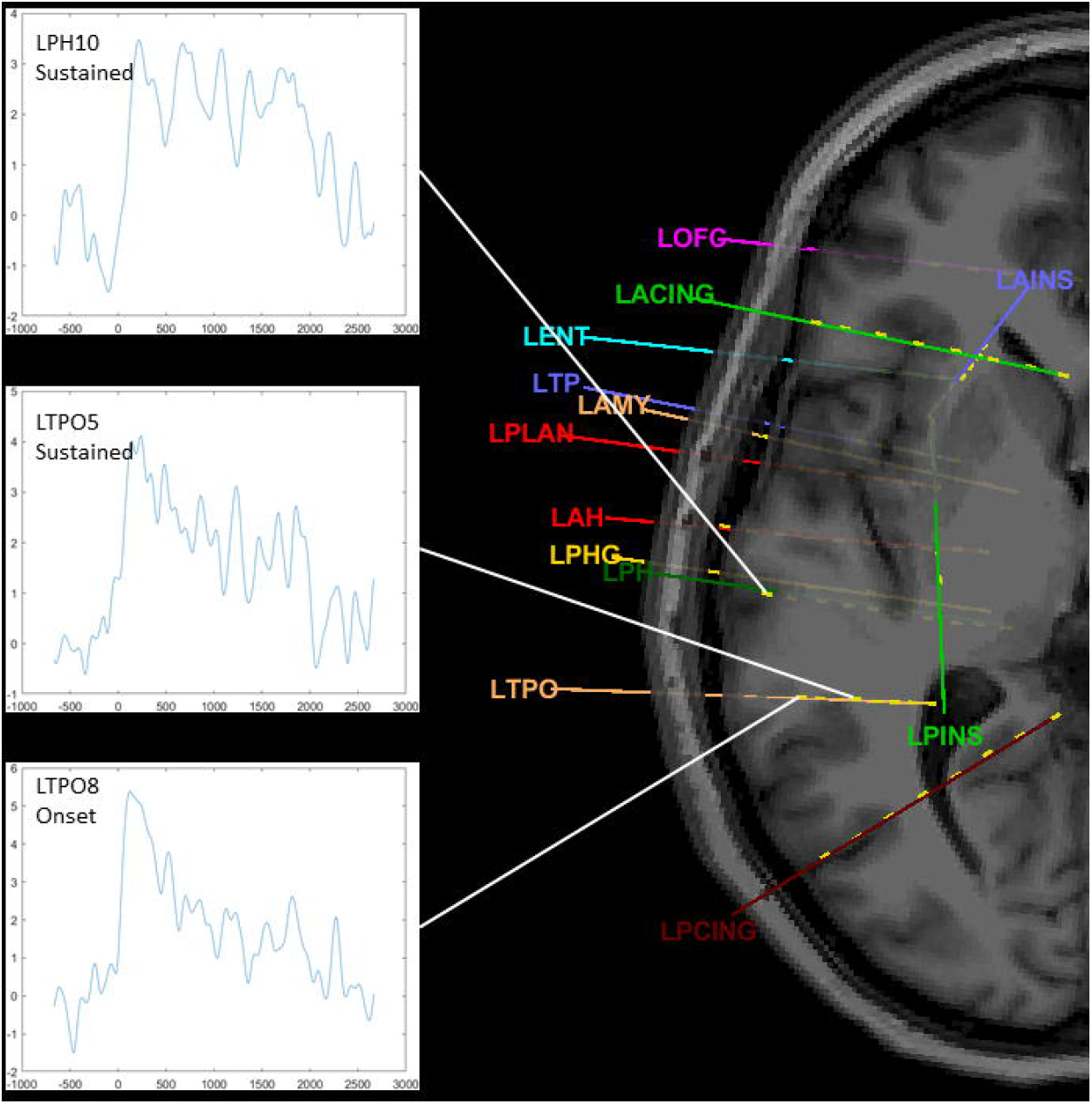
Qualitative Difference between Deep Sustained and Lateral Sustained Sites. Selected representative sites from patient 9 who expressed Deep Sustained, Deep Lateral, and Onset sites. Graphs shown are profiles of average high gamma activity across 70 to 110Hz with y-axis of Arbitrary Units (A.U.) representing values relative to baseline and x-axis representing time in milliseconds (ms). LPH10 is a Lateral Sustained site, LTPO5 is a Deep Sustained site, and LTPO8 is an Onset site; all are related to the superior temporal lobe.

Forseth et al 2020^9^ described that Onset sites of the planum temporale were rarely associated with activity during articulation while Sustained sites frequently were, demonstrating a functional dissociation. Therefore, we performed response-onset analysis to determine whether reactivation occurred during articulation. Across the 100 Sustained sites, reactivation during response articulation occurs more often than chance, 67% of sites active (p_corr_ = 0.010), per right-tailed t-test comparing the rate to 50% chance with Bonferroni correction for 40 statistical comparisons of gamma-augmentation during paired sound stimuli. Across the 24 Onset sites, we did not find the rate of activity to be different from chance, 46% of sites active (p = 0.346), per left-tailed t-test comparing the rate to 50% chance.

Findings related to Alarm and Instrument sounds at Onset and Sustained sites are detailed in Supplement 1. In summary, a plurality of both Onset and Sustained sites of the temporal lobe showed statistically significant high gamma augmentation to both Alarm and Instrument stimuli albeit with insufficient statistical power to demonstrate that this occurred more often than chance. Peak activity at both types of sites under each condition did not differ across sites and patients.

## Discussion

We undertook a specific evaluation of auditory Onset sites of the temporal lobe utilizing iEEG data acquired during an auditory stimulus consisting of paired stimuli of differing perceptual quality but matched fundamental characteristics. As expected, Onset sites reproducibly clustered in the posterior superior temporal region both within and across patients as well as responded to all presented stimulus types when following silence. As predicated by the human voice detector hypothesis^7^, Onset sites demonstrated reactivation in response to a sudden state change of the stimulus. This finding of recurrent auditory Onset site activation in the posterior superior temporal region in the absence of any silent pause is novel. Temporal lobe Auditory Sustained sites also clustered in the superior temporal region but with much less specific localization both within and across patients, generally being situated more rostrally than Onset sites. In contrast to Onset sites, Sustained sites exhibited high gamma augmentation only in response to forward and reverse speech and never to signal correlated noise regardless of position as first or second in a stimulus pair.

Our replication of previous pSTG observations regarding Onset and Sustained sites validates the methodology used in the present study. In addition, the novel finding that Onset sites are capable of reactivation upon abrupt auditory stimulus state change despite lack of a silent pause supports the hypothesis that Onset sites in the pSTG subserve a human voice detector function. Early, transient cortical language processing within the pSTG was not detected in previous studies devoid of intracranial recordings and appears to rely on the technical advantages unique to intracranial electrophysiological measurement of high gamma activity. The present study demonstrates that Sustained sites respond to forward and reverse speech but not to signal correlated noise while Onset sites are less discriminating, at least when following silence. Due to poor sampling in the non-dominant hemisphere, we were unable to replicate the previous finding that both cerebral hemispheres exhibit similar spatio-temporal phenomena of Onset and Sustained high gamma profiles. Utilizing a machine learning algorithm in a model-independent approach, Hamilton et al. 2018^8^ deployed a passive natural speech listening task with superior temporal high gamma activity segregating into profiles representing Onset and Sustained sites. Bilateral Onset sites were more localized and caudal with high temporal and low spectral selectivity, the opposite being true for Sustained sites. Hamilton et al. demonstrated Onset sites to possess shorter latency than Sustained sites, a finding reproduced in our cohort. Forseth et al. 2020 further evaluated Onset and Sustained activity of the superior temporal region utilizing both subdural as well as stereotactic depth electrodes placed in the superior temporal region beneath cortex of the planum temporale^9^. They also found Onset sites localized more posteriorly and superiorly in temporal cortex, with Sustained sites generally more rostral and widespread. Forseth et al. observed that Onset site activity in the planum temporale was uniquely suppressed during articulation, a finding that we reproduced in part by demonstrating that Sustained sites reactivate during articulation more often than chance while Onset sites do not. Thus, our study replicates previous findings regarding the location of superior temporal region auditory Sustained and Onset sites as well as their functional relationships to each other as well as more distant cortical processes.

Similar superior temporal region responses to language related auditory stimuli have been described as part of work that did not utilize the response classifications described here for Sustained and Onset sites. For example, Zheng et al. 2021 report iEEG data using a dynamic analytical approach to high gamma changes following auditory and other stimuli as part of a naming task^15^. They deployed a unique method of parcellating the superior temporal gyrus to elaborate cortical processing in relation to parallel language networks. When they preceded language stimuli by 1s with a warning tone of 500ms duration, superior temporal sites responding to the warning tone were identified in the ‘center’ or ‘middle’ superior temporal gyrus. They hypothesized that this response represented “activation of the primary auditory cortex and auditory beltway areas”^15^, with activations to words being more widespread. Assuming their single word stimuli were less than 1s in duration, Zheng’s procedures would not likely have distinguished Onset from Sustained activation sites. However, the lateral superior temporal sites responding to both non-verbal pure tone warning sounds and then, after a >200ms pause, responding again to the word stimulus were likely Onset type. In a study of temporal cortical responses to dissonant and consonant piano chords, Foo et al. 2016^16^ described sites of the middle to posterior superior temporal gyrus responding with high spatial localization. It is possible that they were also studying Onset sites, as all subdural electrodes with high gamma augmentation in their study were active only during the first half of the 750ms sound stimuli. Foo’s findings suggest that Onset sites present in the non-dominant hemisphere can be further subcategorized into a rostral class more responsive to sounds of increased ‘roughness’ and a caudal class showing no distinction.

Understanding of the role of the posterior superior temporal region in the location and mechanism of auditory language processing is evolving^1,3,4^. Previously, this region has been assumed to subserve phonological processing at or near a cross-roads between the dorsal and ventral language processing streams^4,17^. In human patients, support for such a phonological role is strong across multiple modalities^4^. Repetition errors and phonological paraphasias are best associated with stimulation or lesioning of the posterior superior temporal region^17^. However, the most up-to-date and practical models of language function based on clinical findings fail to distinguish the phenomenon of Onset sites within posterior aspects of the superior temporal region. It is difficult to envision how the transient cortical activity of Onset sites, only briefly responsive to state changes in afferent auditory signals, could directly mediate phonological processing.

Hamilton et al. 2018 aptly described one important challenge to be determining how dividing auditory cortex into Onset and Sustained sites could support multiple levels of auditory processing that is fully integrated at higher cortical levels in support of language comprehension, suggesting that there is a missing component to modern models of language processing^8^. In the current study, we found evidence of human voice detection as a principal functional component of Onset sites. While we noted the novel finding of recurrent Onset activation during a stimulus condition absent a silent pause, we did not find similar recurrent activation in the case of state change from forward speech to SCN. When recurrent activation occurred in this context, its relative intensity depended upon what type of change occurred, implying a more dynamic process. Parallel processing for speech perception is a well-accepted model for neurobiology of language, and Hamilton et al. 2018 make an excellent case that posterior superior temporal Onset sites are likely part of the ventral stream^8^, a bilaterally represented arm of parallel speech processing concerned with the ‘what’ aspect of linguistic information. Detection of a human voice may be part of, but not comprehensive of, the onset detectors of the posterior superior temporal region.

Aiming to maintain a succinct, testable hypothesis describing the underlying function of Onset sites of the posterior superior temporal region, we consider why recurrent Onset activation did not always reproduce that which preceded it. Onset sites responded to any of the three stimulus types when positioned first in a pair. This recapitulates prior data that Onset sites ‘strongly responded to silence followed by sound onset’^8^. Onset sites also respond within clips of natural speech whenever speech followed a pause of 200ms minimum duration^8^. Reverse speech tends to induce stronger high gamma responses at both Onset and Sustained sites throughout the superior temporal region of implanted human patients^6^, when reverse speech is the leading sound. High gamma activity has been linked to saliency of stimuli, as best delineated in the context of a painful stimulus and sensory cortex^18^. Therefore, the elevated high gamma response at both Onset and Sustained sites to leading reverse speech implies that reverse speech is more salient than forward speech. We would propose from our present data that the saliency hierarchy among these stimuli is as follows: reverse speech > forward speech >> SCN >>> silence. However, this was only true for the first stimulus in a pair, and thus the order of auditory signals in a pair did influence the relative intensity of high gamma activity between speech conditions. When SCN was the second sound in a pair following forward speech, Onset sites did not exhibit recurrent high gamma augmentation. When SCN followed reverse speech, however, high gamma augmentation did recur albeit with reduced peak activity compared to the preceding reverse speech. A similar effect occurred when forward speech followed reverse speech, i.e. a recurrent but less intense high gamma augmentation. On the contrary, when SCN was the first sound in a pair, both forward and reverse speech induced recurrent high gamma augmentation at Onset sites with intensity no different than that observed during the preceding SCN, despite the lack of a pause between stimuli. A similar effect occurred whenreverse speech followed forward speech, i.e. a recurrent high gamma augmentation no different than the first.

Taken together, these findings suggest that auditory Onset sites of the posterior superior temporal region function as part of a salience detector subject to inhibitory cortico-cortical feedback loops. The core difference between forward and reverse speech stimuli is the presence or absence, respectively, of extractable linguistic information. Given the nature of their activity, linguistic information is more likely extracted at Sustained sites, rather than at the transient Onset sites. Inhibitory feedback from Sustained sites that actively process linguistic information may explain the relative reduction in intensity of recurrent Onset activity that occurs following forward speech compared to what would be expected if relative saliency were the only determinant. We hypothesize that, in the absence of ongoing auditory linguistic processing at Sustained sites, Onset sites show activity at the onset of a new sound with intensity determined by the saliency of the new sound relative to that of any ongoing sound(s). When Sustained sites are actively processing linguistic information, however, such as during natural forward speech, Onset sites are incompletely suppressed. As a caveat to this hypothesis, the salience detector may possess hysteresis of at least 200ms duration, as outlined by Hamilton et al 2018^8^. It also may be partially suppressed during articulation^9^. These caveats more completely describe presumed parallel feedback loops in which Onset sites monitor for sounds of increased saliency, effectively ‘alerting’ Sustained sites to human voice sounds potentially containing linguistic information, and are then incompletely suppressed during active linguistic information processing or articulation. A neuronetwork simulation conceived from findings of preceding animal studies suggested that activity in Onset sites accelerated the reaction time of Sustained sites^10^. Our hypothesized salience detector may operate in a localized fashion in parallel to the larger network apparatuses, directing them in order to turn their attentional resources to sounds of interest, not unlike a switchboard.

The salience detector hypothesis for posterior superior temporal Onset sites will be testable in future work using auditory ‘roughness’,^16,19^ a perceptual characteristic unique to alarm and dissonant sounds that is associated with stimulus saliency and is measurable^16^. Different sound stimulus types, lacking linguistic information and segregated into categories of high or low roughness, may be presented in pairwise fashion in order to isolate the hypothesized salience detection function of Onset sites. A pause between paired stimuli could be introduced at varying lengths to precisely measure inherent hysteresis. However, as sEEG depth electrodes largely replace subdural grid electrodes for extraoperative intracranial recording^20,21^, it may become more difficult to distinguish and study the separate functions underlying Onset and Sustained sites.

Our results show that deep temporal sites classified as Sustained were enriched with statistically significant high gamma augmentation to SCN when following silence, an unexpected finding. Although this result may indicate that deep sites mediate an inherently different aspect of language processing than lateral sites, alternatively deep sEEG contacts may sample from multiple cortical sites, some mediating Onset activity and some Sustained, blending these signals. Intentional placement of sEEG electrodes in or very near to the cortical area of functional interest would overcome this limitation^9^, but is often neither clinically indicated nor feasible. An alternative approach could involve resolving the discrepancy arising from signal mixing from multiple discontiguous cortical sites by instead using the spatially distributed nature of sEEG eletrodes to specific advantage^22^. Such source localization has been performed using sEEG contacts but primarily only for singular dipoles^23^. To date, only one peer reviewed manuscript has described distributed source modeling of sEEG measurement of physiological high gamma activity^24^. Reliable implementation and validation of similar techniques would enable sEEG as a powerful tool for expanded brain mapping.

We here propose and provide evidence for a salience detector hypothesis for the function of pSTG Onset sites. Consistency of paradigmatic definition and testing protocols will be necessary to achieve further progress in this area and to validate these findings across datasets from diverse research teams and institutions. Updated models of language neurobiology should incorporate these and other novel findings regarding superior temporal region Onset sites. Further evaluation and refinement of this hypothesis will enhance our understanding of key aspects of the neurobiology of language.

## Online Methods

This study was approved by the Institutional Review Board of Oregon Health and Science University (STUDY00015589). All participants provided their written, informed consent/assent prior to testing.

### Participants

The study included participants: (i) with a history of intractable focal epilepsy who were scheduled for extraoperative iEEG recording as part of a presurgical evaluation at Oregon Health and Science University Hospital or Doernbecher Children’s Hospital, Portland, Oregon, between December 2017 and December 2020, (ii) who were aged 5 to 65 years, and (iii) who had acquisition of iEEG signals during a language task as described below. Study exclusion criteria consisted of: (i) presence of massive brain malformations confounding anatomical landmarks for the central sulcus and Sylvian fissure, (ii) history of hearing impairment, (iii) verbal comprehension index (VCI), performance speed index (PSI), Wechsler test of adult reading (WTAR), controlled oral word association test (COWAT), or global intelligence quotient (IQ) with standard score (SS) < 70, (iv) inability to complete the language task described below for any reason, (v) psychiatric history including suicidal ideation, schizophrenia, or bipolar depression type I.

Hemispheric language dominance was determined by findings from clinical fMRI and/or Wada testing, when available. In patients without or having inconclusive evidence from fMRI or Wada testing, hemispheric language dominance was inferred from hand dominance as reported during a formal neuropsychiatric evaluation; i.e. a report of right hand dominance led to the assumption of left hemispheric language dominance.

### iEEG Acquisition

All patients had intracranial electrodes placed; subdural or stereotactic depth, or both, as clinically determined. Only electrodes unaffected by electrical noise or recurrent artifacts were used in subsequent analysis. When subdural electrodes were placed, they consisted of platinum-iridium electrode grids and/or strips embedded within a silastic sheet (PMT Corporation, Chanhassen, MN; 10mm inter-contact distance; 4mm diameter). When stereotactic depth electrodes were placed, they consisted of platinum-iridium electrode contacts (PMT Corporation; 0.8mm diameter, 2.0mm length cylinders; adjacent contacts separate by 0.5-5.25mm). When subdural electrodes were placed, a craniotomy was performed and any ipsilateral stereotactic depth electrodes were placed manually with assistance from a stereotactic cranial navigation system (Brainlab AG, Munich, Germany) utilizing anatomical MRI. When stereotactic depth electrodes were placed in the absence of craniotomy, implants were made utilizing the Robotic Surgical Assistant (ROSA; Zimmer-Biomet, Warsaw, IN) referencing both anatomical MRI and intraoperative CT imaging taken with fiducial screws in place. Each stereotactic depth electrode had 6-16 contacts and multiple were implanted, when utilized. Of the 17 patients, 12 had only stereotactic depth electrodes, 2 had only subdural electrodes, and 3 had a combination of the two.

Extraoperative video-iEEG recordings were obtained for 2-7 days. We deployed the Detroit Procedure for research data acquisition.^25^ A 128-channel Natus Xltek system (Natus Neuro, Middleton, WI, USA) was utilized to acquire signals for all patients except for patient 14 whose data was acquired utilizing a Cadwell Zenith system (Cadwell Industries Inc., Kennewick, WA, USA). Data acquisition was performed at a sampling frequency of 1000Hz. Number and type of electrode contacts are presented in **Table 1** for each patient. The audio line out jack from a sound recorder (Digital Voice Recorder WS-852, Olympus Corporation, Tokyo, Japan for patients 1-12 and Zoom H2n Handy Recorder, Zoom Corporation, Tokyo, Japan for patients 13-17) was wired into iEEG system electrode inputs to record the audio signal from room sounds simultaneous with iEEG signals to optimize synchronization for subsequent analysis.

### Coregistration of iEEG Electrode Contacts on Three-Dimensional MRI

Magnetic resonance imaging (MRI), including volumetric T1- and T2-weighted imaging of the entire head, was obtained pre-operatively per clinical protocols for all patients. Both T1- and T2-weighted images were entered into Freesurfer software (Freesurfer, MGH Harvard, Boston, MA, USA) to reconstruct pial surfaces and delineate cortical anatomy in three-dimensional space. All patients underwent intra-operative or post-operative stereotactic computed tomography (CT) with iEEG electrodes in place. These pre-implantation Freesurfer-reconstructed MRI and post-implantation CT imaging data were imported to Brainstorm software (Brainstorm, University of Southern California, Los Angeles, CA, USA) where iEEG electrode contacts could then be visualized on and within individual brains. For construction of a composite atlas, iEEG contact sites were transformed to ICBM152 space to represent active electrode sites across the cohort. The dominant hemisphere of each patient was mapped to the left hemisphere of the atlas brain; i.e. the dominant right hemisphere of patient 5 was mapped to the atlas’ left hemisphere.

### Task Paradigm

We designed a language task that delivered a consistent set of stimuli with a high likelihood of generating auditory augmentation in high gamma frequency iEEG signal. We generated a total of 63 trials consisting of 7 different stimulus types. The first of these stimulus types involved a set of 9 recorded auditory questions, spoken in native English by author ECB similar to those used previously by Brown et al^6,7^ with an average duration of 2s. The purpose of these question trials was to allow for classification of auditory electrodes similar to Brown et al.^7^ as described below. The remaining 6 stimulus types were 2s total duration pairs of 1s duration stimuli of three to five syllable single words spoken by ECB in either their natural ‘forward’ temporal orientation, temporally flipped ‘reverse’ orientation, or their ‘signal correlated noise’ (SCN) equivalent, similar to that of Brown et al.^7^ Reverse sounds were generated from forward sounds using the *reverse* effect function of Audition software (Adobe Creative Cloud, Adobe Inc., San Jose, CA, USA). SCN sounds were generated from forward sounds utilizing an in-house Matlab (The MathWorks Inc., Natick, MA, USA) script which performs the following processing sequence: 1) performs fast Fourier transform upon the forward speech waveforms, 2) randomizes the phases of all spectral components to destroy spectral information, 3) performs inverse fast Fourier transform to yield speech spectrum noise, 4) applies the amplitude envelope of the original forward speech sound, and 5) scales the maximum amplitude of the resulting SCN to that of the original forward speech sound; producing sounds not unlike those described by Schroeder.^26^ Each pairing involved two different instances of forward, reverse, or SCN with no individual sound being presented more than once. A total of 18 forward sounds were created and subsequently 18 each of reverse and SCN sounds yielding a total of 54 stimulus pair trials. The sound pairings combined with the question stimuli were presented in random order with interstimulus interval varying from 6s to 10s utilizing Presentation software (Neurobehavioral Systems Inc., Berkeley, CA, USA). Patients were instructed to simply listen calmly to the sounds and when they heard a question to simply answer with the first 1- or 2-word noun that came to mind; e.g. Question: “What flies in the sky?” Response: “A bird”.

In addition to the above, patients 13 to 17 underwent an additional 18 trials consisting of 9 each of alarm and instrument sounds. These alarm and instrument sounds were obtained directly from the authors of a previous fMRI study of cortical processing of audible sound ‘roughness’.^19^ These alarm and instrument sounds were matched for duration (either 0.5s or 1s), fundamental frequency, and root mean square power. These 18 trials were presented singularly and integrated into the task in random order with interstimulus interval varying from 6s to 10s. Patient 17, who was bilingual (English and Spanish), additionally received the Spanish language version of the 9 question trials as spoken by the first author (ECB). This was in addition to the default English language version of the questions.

### Digital Signal Processing and Event-Related Analysis

iEEG datasets from the Natus or Cadwell systems were exported in European Data Format for import to EEGLab software (EEGLab, Swartz Center for Computational Neuroscience, Institute for Neural Computation, University of California San Diego, San Diego, CA, USA) for analysis. With aid from visualization and playback on Audition, trial and response onset times were marked. The entire dataset was then low pass filtered at 300Hz followed by high pass filtering at 0.5Hz. The dataset was then referenced to a common average excluding channels with noise, recurrent artifacts, frequent spike-and-wave discharges, or undetectable signal. Each trial was then cropped as an epoch from −1000ms to 3000ms relative to the marked onset and segregated into their respective type. Responses to the question trials were selected for response onset analysis if the response was correct and was inclusive to a collection of responses with response times not varying by more than 2s. Response epochs were similarly cropped starting from 1000ms earlier than the longest response time to 2000ms relative to response onset. All epochs were inspected for evidence of epileptiform spike-and-wave discharge or artifact that effected more than one contact on difference electrodes, the occurrence of which would lead to exclusion of that particular trial epoch. The first 800ms of each epoch, a period of silence, was selected for that trial’s baseline and the baseline’s broad-spectrum power was removed from the epoch. Epochs of the same trial type were then combined to determine event-related spectral perturbation (ERSP). This was performed by decomposition of the signals into time-frequency components through a 3-cycle wavelet method using a Hanning-tapered window. Decomposed data point attributes were adjusted to represent perturbation, augmentation versus attenuation, over 10ms by 5Hz. This ERSP was statistically evaluated at each bin by a bootstrap statistic with a significance level 0.05 after false discovery rate detection for multiple comparisons. As described previously,^7^ additional manual correction was employed by which ERSP values on a given electrode were declared significant only if a minimum of eight time-frequency bins contained within the high gamma range from 70 to 110Hz were arranged in a continuous array spanning (i) at least 20Hz in width and (ii) at least 20ms duration.

### Categorization of Electrodes with Significant High Gamma-Augmentation

Categorization of electrode sites meeting criteria for statistically significant high gamma augmentation was performed entirely from results among the English-language question stimulus trials. As previously described in Brown et al. 2014,^7^ a given electrode is defined as an Auditory site if 1) significant gamma-augmentation begins within 300ms following stimulus onset and 2) ends prior to 300ms following stimulus offset.

Brown et al. 2014 delineated subcategories of Auditory sites labelled Early Auditory and Full Auditory. Here the respective terms Onset and Sustained, similar to Hamilton et al. 2018, are used^8^. Brown et al. 2014 described Onset sites to be those active during the first half of the question stimulus but not the second half. As Onset contacts are the sites of primary interest and our 9 trials have less statistical power than the Brown et al 2014 study, we modified this criterion to minimize the chance of incorrectly classifying a Sustained site as that of Onset. The baseline normalized spectrogram values of the ERSP from 70-110Hz were averaged for each time epoch. Peak value during the first half of the question stimuli from this averaged waveform was determined and compared to that of the second half. An Auditory site is determined to be of the Onset type if the peak during the second half of the question, i.e. beyond 1000ms after stimulus onset, is less than half that during the first. All other Auditory sites were classified as Sustained.

### Statistical Procedures Comparing Outcome Measures Between Trial Types

We asked three distinct categories of questions answerable with statistical tests. First, which sites exhibit statistically significant activity? (Event-related analysis is described above.)

Second, how does activity at Sustained and Onset sites proceed in each of our trial types of primary interest? Two complementary statistical approaches were used here. We determined the latency (L) of statistically significant gamma augmentation from stimulus onset, which then served to guide analysis. 1) At a given active electrode contact, we determined if statistically significant gamma augmentation occurred in each portion of the paired stimulus epochs. 2) We determined peak normalized augmentation, as averaged from 70-110Hz, in each portion of the paired trials. For both approaches, L determined the time boundaries for evaluation. That is, the first portion of the pair was evaluated from time L to 1000ms following stimulus onset with the second portion evaluated from time 1000ms+L to 2000ms+L. After extracting this individual electrode contact level information, we then combined this data across contacts and patients to perform statistical tests. The presence or absence of augmentation during paired trials was determined by a one-sided t-test to test if that portion of the paired trial was active more often than 50% of the time across patients. For peak values, we compared the first portion of the pair to the second portion across patients with a non-parametric Wilcoxon signed ranks test. We then applied Bonferroni correction across all statistical tests contributing to evaluating this question.

Third, how are Onset and Sustained sites temporally related? We selected latency (L) findings from only those patients who contributed both Onset and Sustained electrode sites. We then performed a non-parametric Wilcoxon rank sum test to determine if Onset and Sustained sites have different L values.

The above-described statistical tests were planned. All other statistical tests were therefore unplanned and strict Bonferroni correction was broadly applied to them accounting for all preceding statistical tests regardless of which question they aimed to answer. Throughout, the alpha threshold is 0.05. Corrected *p-values* for a statistical test are represented as p_corr_, with ‘p’ indicating an uncorrected value.

## Supporting information

Supplement Results

## Acknowledgments

We acknowledge patients and families for their interest and willingness to participate in this study as they journeyed on their individual paths toward improved seizure control and quality of life. We very much appreciate the Oregon Health and Science University’s Clinical Neurophysiology team of the Department of Neurology for assistance along the way, especially Aaron Kawamoto, Allison Gardner, Cecily Key, Emily Stair, Inrun Kaur, Kevin Vinecore, and the team’s Director Ilker Yaylali, MD, PhD. The OHSU Adult and Doernbecher Pediatric Epilepsy Teams were helpful in guiding patients to feel comfortable with the study, especially Jason Coryell, MD, Colin M. Roberts, MD, Carter D. Wray, MD, Ittai Bushlin, MD, PhD, Paul V. Motika, MD, Lia Ernst, MD, and David Spencer, MD. The help and patience of the nursing staff of the Pediatric Intensive Care Unit of Doernbecher Children’s hospital as well as the Neurosciences Intensive Care Unit and the Epilepsy Monitoring Unit of OHSU Hospital were critical to data collection and patient comfort. The contribution of David Dunham of OHSU’s Clinical Technology Services for providing necessary tools and skills for creation of custom hardware to integrate research audio equipment with clinical EEG acquisition hardware. We thank Shirley McCartney, PhD, for editorial guidance from conception to completion and OHSU Department of Neurological Surgery’s research coordinators especially James “Obi” Obayashi and SamanthaYau for IRB regulatory and compliance assistance. Multiple other individuals aided in some form along the way including Paxton Gehling, MD, Karan Rai, Kelly Collins, MD, and others and we are so grateful for their interest and support. Without all the hard-working, helpful people we are surrounded by, we would not be able to see such a project to completion. Finally, the first author is forever grateful to Eishi Asano, MD, PhD, of Children’s Hospital of Michigan, who has always taken the concept of mentorship to heart and follows through with the concept that it can and should be lifelong.

## Author Contributions

The first and corresponding author ECB conceived of, developed, constructed data acquisition devices, assembled stimulus delivery system, performed data acquisition of iEEG signals, analyzed all electrophysiological signals, contributed to image processing, interpreted results, and constructed the initial draft of this manuscript along with contributing to all stages of editing. The co-author BS assisted with data acquisition, performed image processing especially involving localization and representation of iEEG electrode contact positions in three dimensional space in both individual as well as atlas space, provided insight into results interpretation, and contributed to all stages of editing of this manuscript. The co-authors AMR and NRS led all integration between clinical and research work, provided insight into results interpretation, and contributed to all stages of editing of this manuscript.

