## Supplement Results for "Speech-related auditory salience detection in the posterior superior temporal region"

**Cortical Responses of Onset and Sustained Temporal Auditory sites to Alarm and Instrument Sounds.** Alarm sounds have been described to possess a unique audible sound characteristic known as ‘roughness’ which is not present in non-alarm sounds. It is thought that our brains respond preferentially to perceived roughness.^1,2^ It has been shown by iEEG that the posterior superior temporal gyrus is involved in this activity.^2^ As part of an unplanned preliminary evaluation of this phenomenon, we included matched Alarm and Instrument sounds obtained from Dr. Arnal^1^ as isolated and randomly presented trial stimuli for patients 13 through 17. Among these patients’ sites there were 6 identified as Onset and 28 identified as Sustained in the temporal lobe. Among Onset sites, 5 sites (83%) showed high gamma augmentation to both Alarm and instrument sounds; statistical tests were not performed to compare to 50% chance due to anticipated low statistical power. These Onset sites did not show a significant difference between peak ERSP associated with Alarm and Instrument conditions; p-value 1.0 per Wilcoxon signed rank test. Among Sustained sites, 75% showed high gamma augmentation to Alarm sounds while 71% showed high gamma augmentation to Instrument sounds, neither being elevated above the statistical effect of chance; p_corr_-value 0.1276 and p_corr_-value 0.4488, respectively, per right-sided t-test comparing the rate to 50% chance and corrected by Bonferroni method for 44 statistical comparisons. These Sustained sites did not show a significant difference between peak ERSP associated with Alarm and Instrument conditions; p-value 0.509 per Wilcoxon signed rank test. **Figure S1** displays representative Onset and Sustained sites responding to Alarm and Instrument stimuli; qualitatively, the activity related to each is similar.

**Figure S1 – Cortical Response of Temporal Onset and Sustained sites to Alarm and Instrument Stimuli**

Depicted are representative Onset and Sustained sites responding to Alarm and Instrument sounds. All selected sites reached statistical significance for high gamma augmentation. Graphs shown are profiles of averaged high gamma activity across 70 to 110Hz with y-axis of Arbitrary Units (A.U) representing values relative to baseline and x-axis as time in milliseconds (ms).

**Video S1 – Cohort Level Representation of All Active Temporal Lobe Auditory Sites**

Supplementary video of the cohort level representation of sites displayed on the atlas brain as in Figure 2 turning first horizontally and then rolling to aid appreciation electrode locations in three-dimensional space. Left atlas hemisphere represents the language dominant hemisphere.
