## Supplementary figures and images for "Speech-related auditory salience detection in the posterior superior temporal region"

### Supplement Figure S1

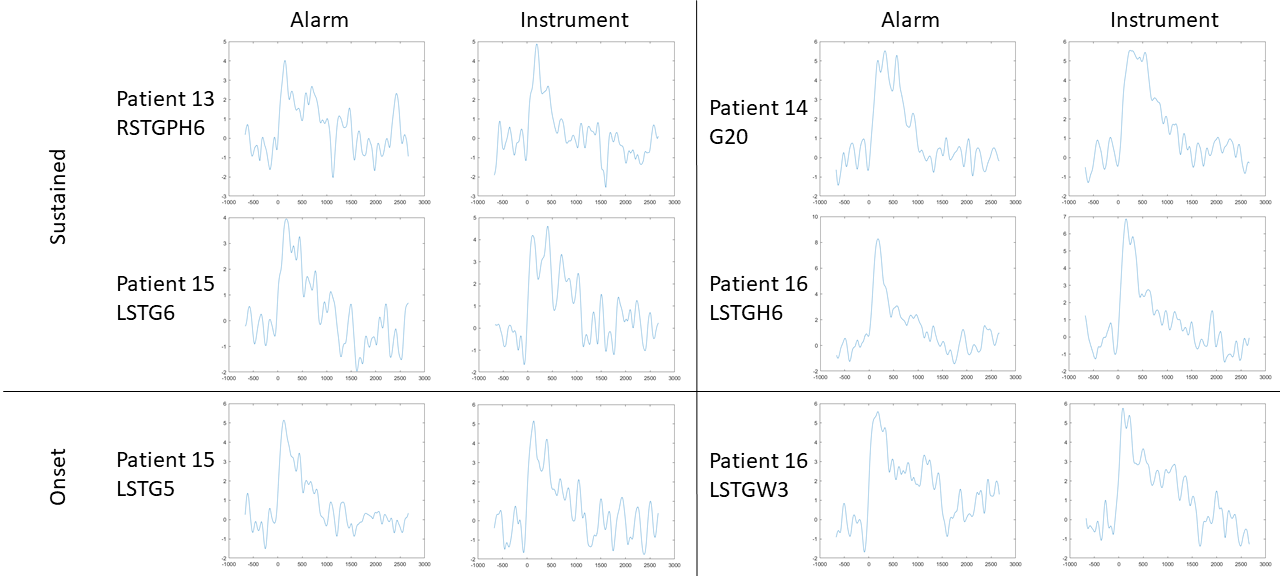
